# Restoration of locus coeruleus noradrenergic transmission during sleep

**DOI:** 10.1101/2024.07.03.601820

**Authors:** Jiao Sima, Yuchen Zhang, Declan Farriday, Andy Young-Eon Ahn, Eduardo Ramirez Lopez, Chennan Jin, Jade Harrell, Dana Darmohray, Daniel Silverman, Yang Dan

## Abstract

Sleep is indispensable for health and wellbeing, but its basic function remains elusive. The locus coeruleus (LC) powerfully promotes arousal by releasing noradrenaline. We found that noradrenaline transmission is reduced by prolonged wakefulness and restored during sleep. Fiber-photometry imaging of noradrenaline using its biosensor showed that its release evoked by optogenetic LC neuron activation was strongly attenuated by three hours of sleep deprivation and restored during subsequent sleep. This is accompanied by the reduction and recovery of the wake-promoting effect of the LC neurons. The reduction of both LC evoked noradrenaline release and wake-inducing potency is activity dependent, and the rate of noradrenaline transmission recovery depends on mammalian target of rapamycin (mTOR) signaling. The decline and recovery of noradrenaline transmission also occur in spontaneous sleep-wake cycles on a timescale of minutes. Together, these results reveal an essential role of sleep in restoring transmission of a key arousal-promoting neuromodulator.

## Main

Sleep is an essential innate behavior indispensable for maintaining optimal brain function^1^. Acute sleep deprivation can lead to reduced alertness and impairment of a wide array of cognitive functions, including attention, learning, and memory consolidation ^2^, which can be reversed by recovery sleep. However, the neural mechanisms underlying the restorative effects of sleep remain elusive.

Neurons in the locus coeruleus (LC), part of the ascending arousal system ^3^, play a powerful role in promoting wakefulness and arousal ^4–10^ and regulating cognition ^11^. LC neurons release noradrenaline (also called norepinephrine, NE) through their brain-wide projections ^12^. NE signaling is essential to the wake-promoting effect of LC neurons, as CRISPR-based knockdown of dopamine beta-hydroxylase (DBH), which catalyzes the synthesis of NE from dopamine, reduced their wake-promoting effect ^13^.

LC neurons exhibit substantially higher activity during wakefulness than rapid eye movement (REM) and non-REM (NREM) sleep ^14–16^. Previous studies suggest that prolonged LC activation or sleep deprivation can deplete NE ^4,17,18^, which could lead to impaired arousal and cognitive functions. However, the magnitude, time course, and mechanism of the reduction and restoration of NE transmission remain unclear. In this study, by measuring NE release using its biosensor, we characterized the effects of sleep and wakefulness, LC neuron activity, and mTOR signaling on the efficacy of LC-NE transmission.

### LC-NE transmission is diminished by sleep deprivation and restored in sleep

We injected adeno-associated virus (AAV) with Cre-dependent expression of ChrimsonR (hSyn-FLEX-ChrimsonR-tdTomato) ^19^ into the LC and AAV expressing a genetically encoded NE sensor (GRAB_NE3.1_) ^20^ into both the LC and medial prefrontal cortex (mPFC) of *DBH-Cre* mice (Fig. 1a,b). Laser stimulation in the LC (635 nm, 4 mW, 50 ms/pulse; inter-pulse interval: 40 – 80 s) evoked robust NE release in both the LC and mPFC, as shown by the transient increases in GRAB_NE3.1_ fluorescence measured by fiber-photometry; the mutant NE sensor showed no laser-evoked response (Extended Data Fig. 1a).

**Fig. 1.**
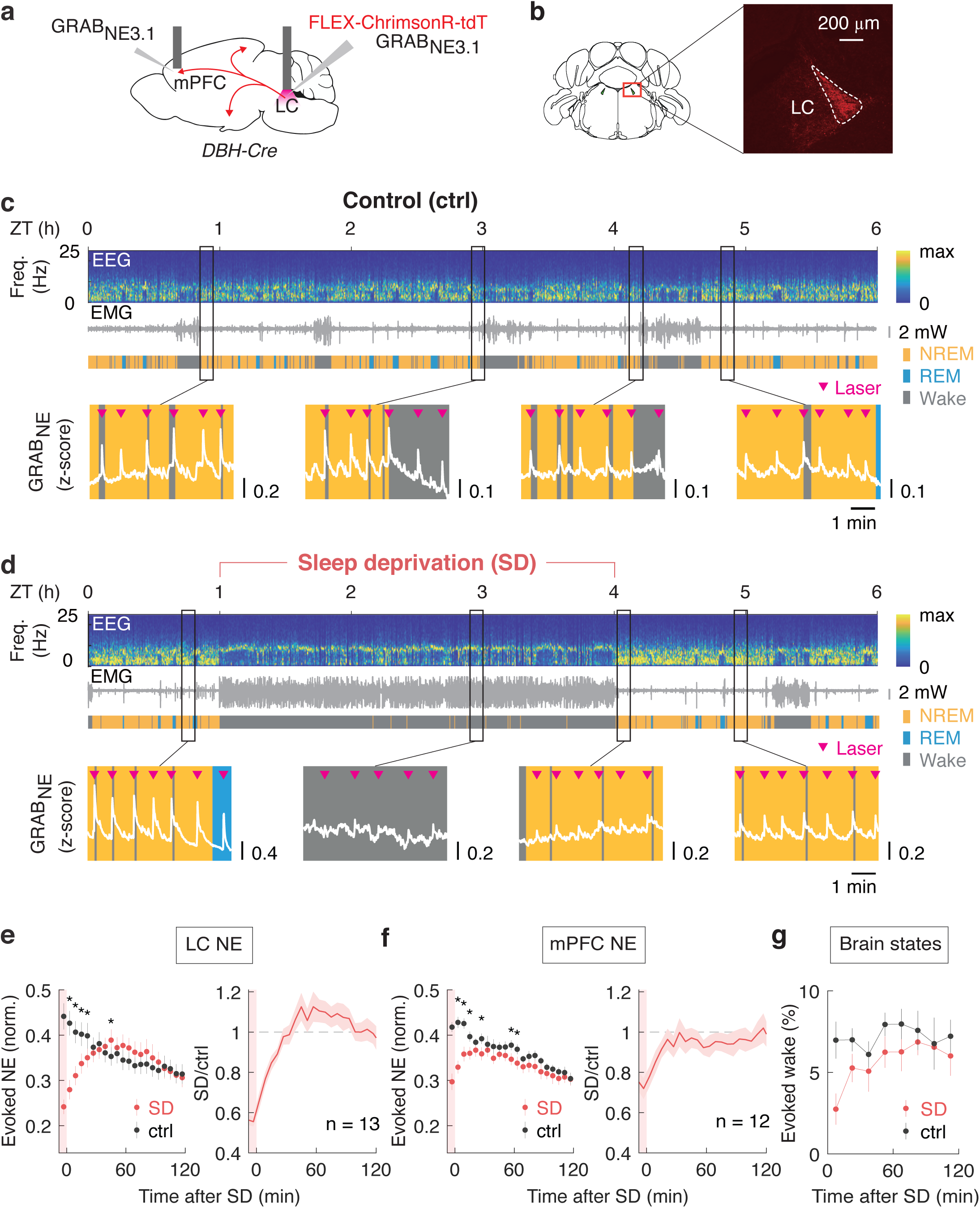
LC-NE transmission is reduced by sleep deprivation and restored during sleep. **a,** Schematic for measuring evoked NE release in LC and mPFC. **b,** Fluorescence image showing ChrimsonR-tdTomato expression in LC-NE neurons. **c,** Example control session showing EEG, EMG, and color-coded brain states. GRAB_NE3.1_ fluorescence (z-score) during selected periods (vertical boxes) are shown on expanded timescale. Pink arrowheads indicate laser pulses. **d,** Example SD session (SD: ZT 1-4) with GRAB_NE3.1_ traces shown at similar time points as in control. **e&f,** Left, laser-evoked NE response for control (black) and SD (red) mice binned every 6 min and averaged across animals in post-SD period in LC (**e)** and mPFC (**f)**. Error bar, SEM. Right, SD/ctrl ratio averaged across all mice. Shading, SEM. P_LC-NE_ < 2.2 × 10^−16^, P_treatment (LC-NE)_ = 0.03, P_time (LC-NE)_ = 1.3 × 10^−13^; P_mPFC-NE_ = 0.009; P_treatment (mPFC-NE)_ = 1.7 × 10^−13^, P_time (mPFC-NE)_ < 2.2 × 10^−16^, * P < 0.05 (two-way ANOVA with Bonferroni correction). **g,** Probability of wakefulness induced by each 50-ms laser pulse binned every 15 min in post-SD period in control (black) and SD (red) condition (n = 13). Error bar, SEM. P_treatment_ = 1.2 × 10^−4^, P_time_ = 0.04 (two-way ANOVA).

After 1 h of baseline measurement (ZT 0-1), we subjected the mouse to 3 h of sleep deprivation (SD), followed by 2 h of recovery sleep (Extended Data Fig. 1b). Laser-evoked NE release was measured continuously throughout each 6 h recording session. Compared to control sessions (‘ctrl’, Fig. 1c), laser-evoked NE release in the LC was markedly reduced by 3 h of SD (Fig. 1d). Immediately after SD, the amplitude of NE release was only ∼60% of the control. However, evoked NE release recovered gradually over ∼30 min of recovery sleep (Fig. 1e, n = 13). Similar SD-induced reduction and subsequent recovery of evoked NE release were also observed in the mPFC (Fig. 1f, n = 12).

We next analyzed the behavioral consequence of the change in NE transmission. The single pulse of optogenetic LC stimulation used for measuring NE release (50 ms) also induced a transient increase in wakefulness as measured by electroencephalogram (EEG) and electromyography (EMG) (Extended Data Fig. 1c,d). Laser-induced wakefulness, measured by the difference in the probability of wakefulness within 20 s before and after the laser pulse (Extended Data Fig. 1e), was significantly lower than the control immediately after 3 h of SD (Fig. 1g), but it recovered gradually during recovery sleep. Thus, SD-induced reduction of NE transmission is associated with an impaired ability of LC neurons to promote wakefulness.

### NE transmission is weakened by prolonged LC activation

Since LC neurons are more active during wakefulness than NREM or REM sleep ^14–16^, the SD-induced reduction of NE transmission could result from prolonged LC activation. To test this possibility, we replaced the SD with optogenetic activation of LC neurons at 0.2 Hz (50 ms/pulse) (Fig. 2a). Immediately after 3 h of LC activation, the amplitude of NE release was reduced to ∼60% of the control, but it recovered over the next ∼30 min (Fig. 2b,c). The wake-promoting effect of LC neurons also showed corresponding reduction and recovery (Fig. 2d). Thus, prolonged LC activation also diminishes NE transmission and the wake-inducing potency of LC neurons, similar to the effects of SD.

**Fig. 2.**
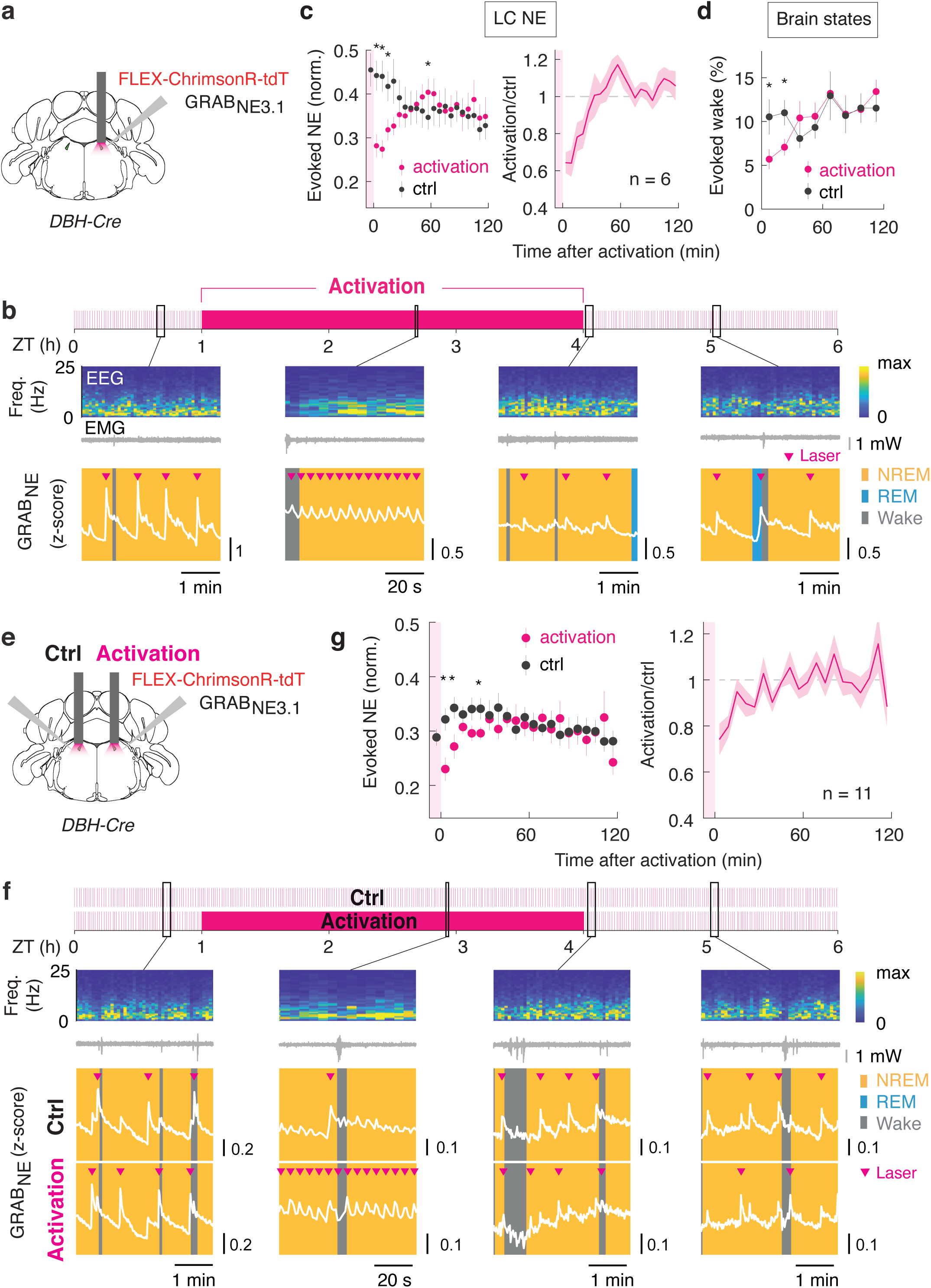
Prolonged LC-NE neuron activation causes reduction of NE transmission. **a,** Schematic for 3-h 0.2-Hz LC activation experiment. **b,** Example LC activation session showing EEG, EMG, color-coded brain states, and GRAB_NE3.1_ fluorescence at selected time points before, during or after 0.2 Hz activation (ZT 1-4). **c,** Left, laser-evoked NE release binned every 6 min and averaged across animals in post-activation period in LC. Error bar, SEM. Right, activation/ctrl ratio averaged across all mice. Shading, SEM. P = 4.0 × 10^−14^, P_treatment_ = 0.8 × 10^−3^ (two-way ANOVA); **d,** Probability of wakefulness induced by each 50-ms laser pulse after 3 h LC activation (red) and in control (black), binned every 15 min (n = 6). Error bar, SEM. P = 0.02, P_time_ = 2.9 × 10^−4^ (two-way ANOVA); **e,** Schematic for LC activation experiment comparing evoked NE release on the activated side to contralateral control side. **f,** Example session showing EEG, EMG, color-coded brain states, and GRAB_NE3.1_ fluorescence at selected time points. Upper, ctrl LC; lower, activated LC. **g,** Left, laser-evoked NE response in post-activation period. Error bar, SEM. Right, ratio between activated and control side averaged across all mice. Shading, SEM. P = 0.006, P_treatment_ = 0.006 (two-way ANOVA); * P < 0.05 (two-way ANOVA with Bonferroni correction).

In addition to activating the LC neurons near the optic fiber, optogenetic stimulation may also affect many other neurons, astrocytes, and microglia through the released NE, which could indirectly affect LC-NE transmission. To control for the effect of global changes in the brain, we compared NE release from the LC neurons subjected to 3 h of activation to that from the contralateral LC, which was much less activated (Fig. 2e, and Extended Data Fig. 2a). Laser-evoked NE release at the activated side was significantly reduced compared to the contralateral side immediately after the 3 h activation, but it recovered over the next ∼30 min (Fig. 2f,g). This further supports a direct role of prolonged LC activation in inducing the decline in NE transmission.

### Genetic inactivation of mTOR signaling slows down recovery of LC-NE transmission

What mechanism underlies sleep-dependent recovery of LC-NE transmission? In addition to reduced release that may help conserve intracellular NE stores, sleep is also known to facilitate cellular repair and rejuvenation by promoting mTOR activity and protein synthesis ^21–23^. To test the role of mTOR signaling in sleep-dependent restoration of LC-NE transmission, we deleted *Rptor* (encoding an obligatory component of mTOR complex 1) specifically in LC-NE neurons by crossing *DBH-Cre* mice with floxed *Rptor* mice ^24^. Compared to wild type (WT) mice, *Rptor*-/-mice showed lower levels of tyrosine hydroxylase (TH) protein, the rate-limiting enzyme for NE synthesis (Fig. 3a,b and Extended Data Fig. 3a,b), similar to the effect of *Rptor* knockout in midbrain dopaminergic neurons ^25^.

**Fig. 3.**
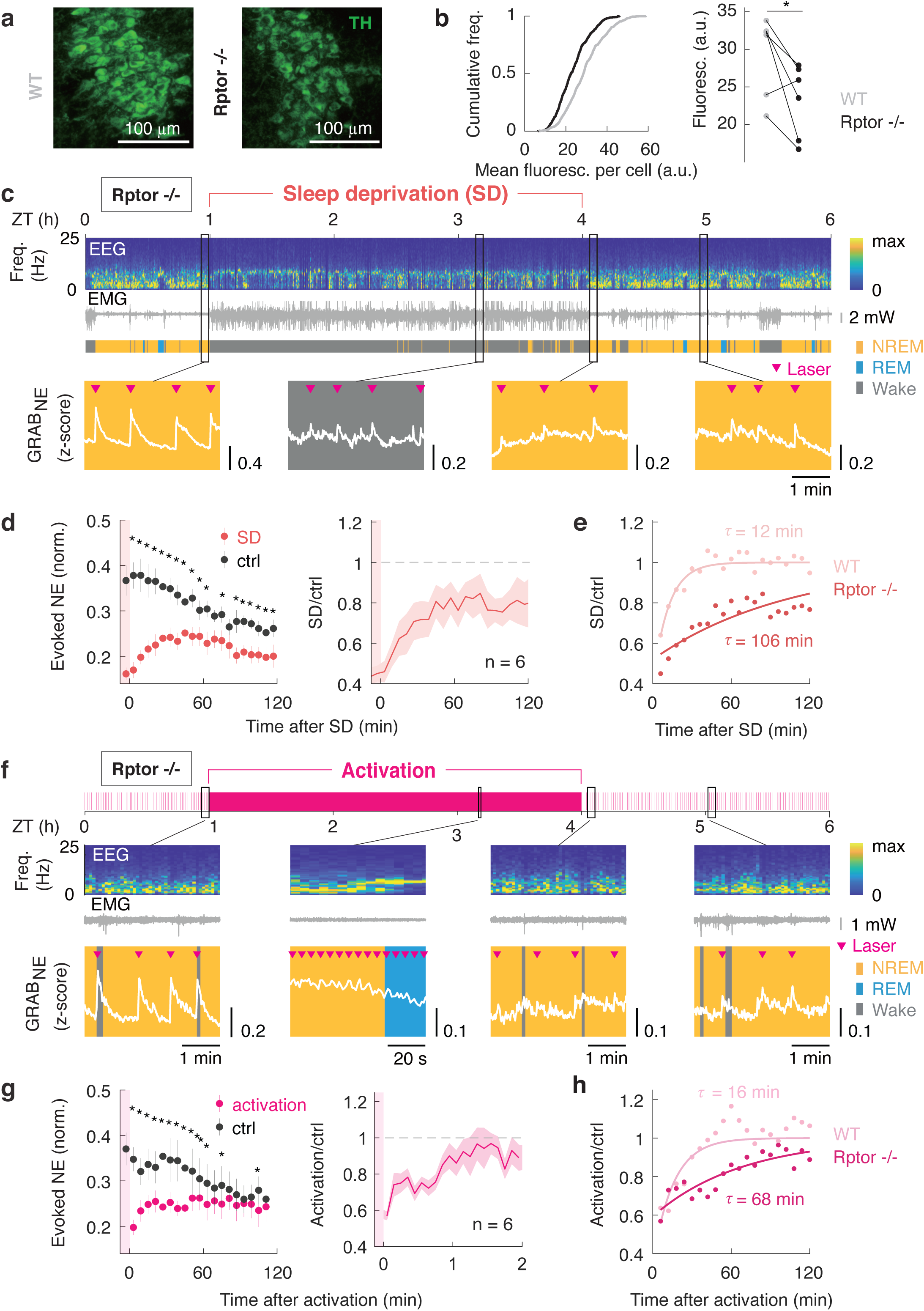
Genetic inactivation of mTOR signaling slows down recovery of LC-NE transmission. **a,** Example images for TH immunostaining (green) in LC in WT and *Rptor*-/-mice. **b,** Cumulative frequency for TH fluorescence in individual cells in *Rptor*-/-(black, n = 6) and WT (grey, n = 6) mice. P = 0.04 (paired t-test). **c,** Example SD session showing EEG, EMG, and GRAB_NE3.1_ fluorescence at selected time points in *Rptor* -/-mutant. **d,** Left, laser-evoked NE response in post-SD period in *Rptor*-/-mutants. Error bar, SEM. P = 2.4 × 10^−5^, P_treatment_ < 2.2 × 10^−16^, P_time_ = 1.9 × 10^−6^ (two-way ANOVA). Right, SD/ctrl ratio averaged across all mice. Shading, SEM. **e,** Single exponential curve fit for laser-evoked NE release after SD for *Rptor*-/- (dark red, n = 6) and WT (light red, n = 6) mice. P_treatment_ < 2.2 × 10^−16^, P_time_ = 1.4 × 10^−9^ (two-way ANOVA). **f,** Example LC activation session for *Rptor*-/-mutant. **g,** Left, laser-evoked NE release in post-activation period for *Rptor*-/-mice. P = 1.7 × 10^−6^, P_treatment_ < 2.2 × 10^−16^, P_time_ = 0.002 (two-way ANOVA). Right, activation/ctrl ratio averaged across all mice. Shading, SEM. **h,** Single exponential curve for laser-evoked NE responses after LC activation in *Rptor*-/- (dark pink, n = 6) and WT (light pink, n = 6) mice. P = 0.01, P_treatment_ = 4.9 × 10^−14^, P_time_ < 2.2 × 10^−16^. * P < 0.05 (two-way ANOVA with Bonferroni correction).

We then repeated the 3 h SD or optogenetic activation experiment in *Rptor*-/- mice (Extended Data Fig. 4). Compared to WT control, *Rptor*-/- mice showed much slower recovery of NE transmission following 3 h of either SD or optogenetic activation, with the time constant of recovery increasing from 10-20 min to 1-2 h (Fig. 3c-h). Thus, the rate of sleep-dependent recovery of NE transmission depends strongly on mTOR signaling in LC neurons.

### Fatigue and recovery of LC-NE transmission in natural sleep-wake cycles

Although the function of sleep is often probed experimentally by several hours of sleep deprivation, in mice the duration of each bout of spontaneous wakefulness is only a few minutes on average ^26^. We asked whether the wake-associated decline and sleep-dependent recovery of NE transmission also operate on the timescale of minutes. Analysis of evoked NE release across spontaneous sleep-wake cycles showed that the release in the LC decreased over several minutes from wake bout onset (Fig. 4a,b) and recovered after mice transitioned into NREM sleep (Fig. 4a, and c left panel). We then divided all the wake bouts into short and long groups. We found that following long wake bouts (> 55 s, median duration of wake bouts), NE transmission was reduced to a lower level, and the recovery to baseline was significantly slower than following short bouts (≤ 55 s; Fig. 4c right panel). Thus, fatigue and recovery of NE transmission also operate in natural sleep-wake cycles on the order of minutes, and the time course of recovery scales with the duration of prior wakefulness.

**Fig. 4.**
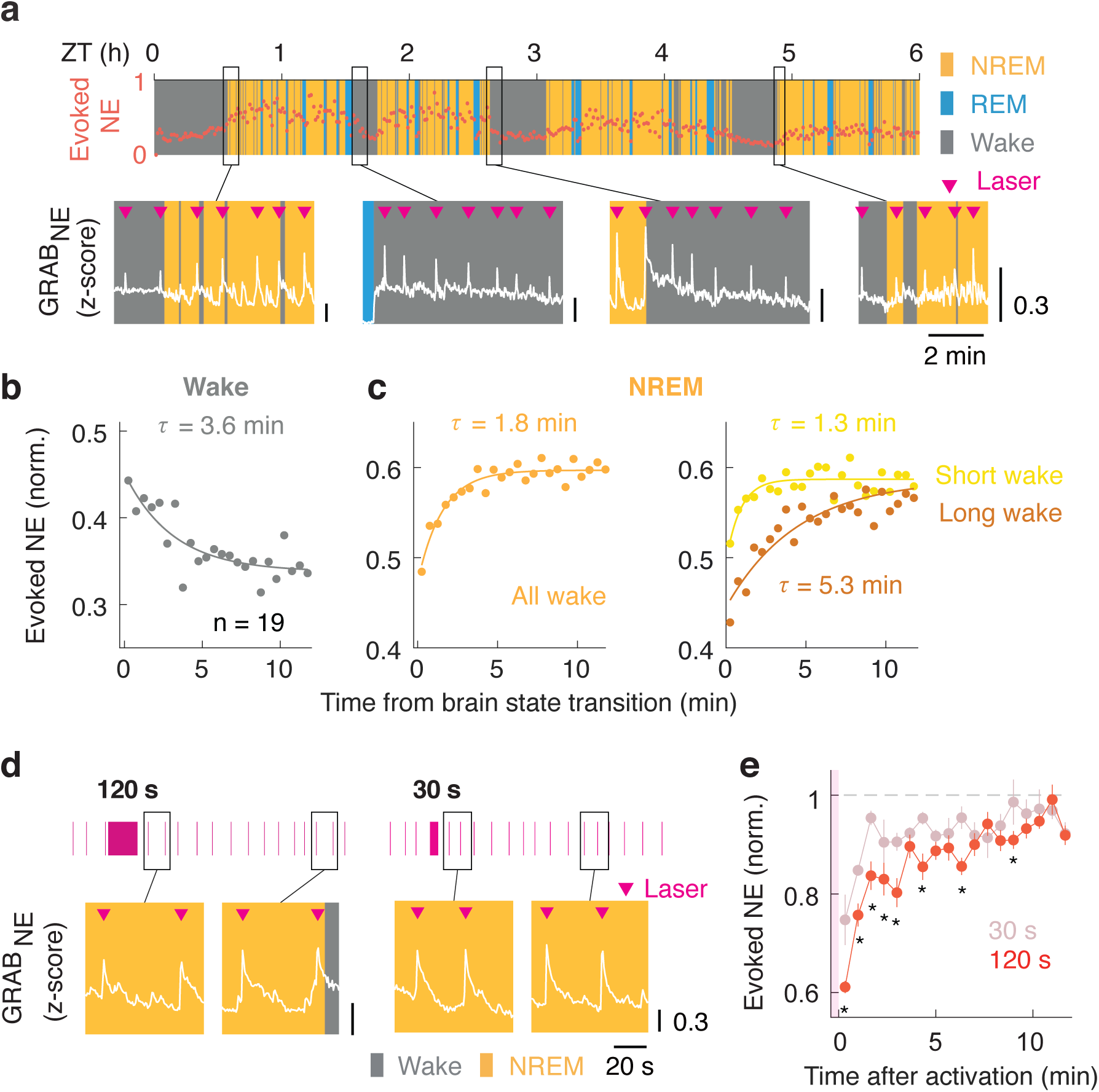
Decline and recovery of LC-NE transmission in natural sleep-wake cycles. **a,** Example GRAB_NE3.1_ fluorescence showing the dynamics of laser-evoked NE response in the LC in natural sleep/wake cycles. **b,** Single exponential curve fit for evoked NE response in the LC after the onset (time 0) of wake bouts. Data points are averaged across trials and binned every 30 s. **c,** Left, single exponential curve fit for evoked NE response in the LC in NREM after wake to NREM transition (time 0). Right, single exponential fits for NREM following short (yellow) or long (brown) wake bouts. P_treatment_ < 2 × 10^−16^, P_time_ < 2 × 10^−16^ (two-way ANOVA). **d,** Example GRAB_NE3.1_ fluorescence showing evoked NE release in LC after 1 Hz LC activation for either 120 s or 30 s. Vertical bars represent 50-ms laser pulses. **e,** Laser-evoked NE response in NREM in LC after LC activation for 30 s (light red, n = 7) or 120 s (red, n = 5). Data points are averaged across animals and binned every 40 s. Error bar, SEM. P = 0.03, P_treatment_ = 2.2 × 10^−9^, P_time_ < 2 × 10^−16^; * P < 0.05 (two-way ANOVA with Bonferroni correction).

The average LC neuron firing rate during quiet wakefulness is ∼1 spike/s ^14,27^. We next applied 1 Hz activation of LC-NE neurons for either 30 or 120 s, comparable to the durations of short and long bouts of spontaneous wakefulness, respectively. Compared to 30 s, 120 s activation caused a stronger reduction of NE transmission and slower recovery (Fig. 4d,e). LC-NE fatigue can also be induced by 120 s of 0.2 Hz activation, but the recovery was faster than that following 1 Hz stimulation (Extended Data Fig. 5).

## Discussion

Sleep is known to provide multiple benefits for the body and mind, ranging from enhancing memory consolidation ^16,28,29^ to regulating immunity ^30^. Here we show that sleep also restores transmission of a key wake-promoting neuromodulator. A previous study based on microdialysis showed that over a 6 h period of SD the NE level in the mPFC declined during the last 1-2 h ^18^. However, it is unclear whether the observed NE reduction was caused by lowered LC neuron firing rates or reduced efficacy of NE transmission. In this study, we measured NE release evoked by a single pulse of optogenetic LC stimulation, enabled by the high temporal resolution of GRAB sensor imaging ^20^. We showed that the efficacy of evoked NE release decreases over the duration of wakefulness and is restored by sleep (Fig. 1).

The reduction of evoked NE release is activity dependent. Even 30 s of stimulation caused a strong reduction (Fig. 4d,e), consistent with previous observations that strong LC activation rapidly reduced NE release ^4,17^. Interestingly, the time course of recovery scales with the duration of wakefulness or LC neuron activation, from minutes (Fig. 4c,e) to hours (Figs. 1, 2). Thus, activity-dependent functional fatigue and recovery of LC-NE neurons can occur on multiple timescales ^17^.

Sleep-dependent recovery of NE transmission (Fig. 1e,f, and Fig. 4c) is likely due, at least in part, to reduced LC neuron activity ^14–16^, which may restore the intracellular NE level by rebalancing the rates of release, reuptake, and synthesis. On the timescale of hours, the rate of recovery of NE transmission depends on mTOR signaling (Fig. 3e,h). The slower recovery in *Rptor*-/- mice could be partly explained by the reduced TH in LC-NE neurons (Fig. 3a,b), which may reduce the rate of NE synthesis. Sleep has also been shown to promote mTOR activity and protein synthesis in general ^21–23,31^. It would be interesting to examine whether other mTOR-dependent mechanisms also contribute to the recovery of LC-NE transmission during sleep.

Our observation that a single pulse of LC stimulation can evoke transient wakefulness (Extended Data Fig. 1c,d) is consistent with the well-known arousal-promoting function of these neurons ^4–9^. The SD-induced decrease in NE transmission diminished the wake-promoting potency of LC neurons (Fig. 1g and 2d), which could contribute to the decreased arousability of the animal after sleep deprivation ^32^, a hallmark of sleep homeostasis ^33^. Given the importance of NE in regulating a broad range of cognitive functions ^11^, its reduced transmission is also likely to contribute to SD-induced cognitive impairment ^2,18^. Moreover, LC neurons are particularly prone to neurodegeneration in both Alzheimer’s and Parkinson’s diseases ^34,35^, partly due to their vulnerability to activity-dependent metabolic stress ^36^. Thus, the restorative effect of their inactivation during sleep is likely to extend beyond the recovery of NE transmission and contribute to the long-term health of LC neurons.

## Acknowledgments

We thank A. Kumar for assisting with histology; N. Dolensek, C. Ma, X. Ding, L. Lu, C. Chen, C. Tso, R. Zoncu, X. Gu, Y. Chen, P. Kosillo and T. Qiu for helpful discussions; H. Bateup for her generosity in sharing the *Rptor* mouse line.

## Funding

This work was supported by the Howard Hughes Medical Institute, National Institutes of Health, National Institute of Neurological Disorders and Stroke grant U01NS113358 (to YD); and the Damon Runyon Fellowship Award DRG-2414-20 (to JS).

## Author contributions

JS and YD conceived the project and experimental design; JS performed majority of the experiments and data analysis; YZ, AYA, ERL, CJ contributed to preliminary results leading to the development of the project; DF carried out the IHC experiment, JH corrected sleep scoring, DD assisted with fiber photometry and statistical analysis; DS set up the sleep deprivation platform. JS and YD wrote the manuscript with input from all authors.

## Competing interests

The authors declare no competing interests.

## Materials request & Correspondence

should be addressed to Yang Dan.

## Data and materials availability

All data are available upon request.

## Methods

### Animals

All experimental procedures were carried out in accordance with the protocol approved by the animal care and use committee at the University of California, Berkeley. Animals were housed on a 12-h dark/12-h light cycle with light onset at 7 am. Adult (>2 months old) male and females were used for all experiments. Mice were typically housed in groups of up to 5 animals. After surgery, animals were individually housed to prevent damage to the implant. Experiments were conducted at least 2 weeks after surgery.

### Breeding strategy

*DBH-Cre* mice (B6.FVB(Cg)-Tg(DBH-Cre)KH212Gsat/Mmucd, MMRRC: 036778-UCD) were crossed with black 6 females (C57BL/6) to produce *DBH-Cre* experimental mice. To generate LC-specific *Rptor* knock-out, male and female mice heterozygous for *Rptor* (Jax 013188) and *DBH-Cre* were crossed (*Rptor^wt/fl^;DBH-Cre^wt/+^* × *Rptor^wt/fl^;DBH-Cre^wt/+^*) to produce experimental animals: homozygous *Rptor KO* (*Rptor^fl/fl^;DBH-Cre^wt/+^*, referred to as *Rptor* -/-) *or WT* (*Rptor^wt/wt^;DBH-Cre^wt/+^*, referred to as WT). For experiments involving comparisons between the two genotypes, age- and sex-controlled mice from the same breeding pairs were used (Fig. 3e,h).

Genotyping primers for DBH-Cre mice are as follows:

Forward: AATGGCAGAGTGGGGTTGGG
Reverse: CGGCAAACGGACAGAAGCATT

### Surgical procedures

Adult mice (6 - 12 weeks old) were anesthetized with isoflurane (5% induction, 1.5% maintenance) and placed on a stereotaxic frame. Buprenorphine (0.1 mg/kg, subcutaneous) and meloxicam (10 mg/kg, subcutaneous) were injected before surgery. Lidocaine (0.5%, 0.1 mL, subcutaneous) was injected near the target incision site. Body temperature was stably maintained throughout the procedure using a heating pad. After sterilization with ethanol and betadine, the skin was incised to expose the skull, and a patch of scalp and connective tissue were removed. Surgeries typically consisted of virus injection followed by optical fiber and EEG/EMG electrodes implantation.

### Virus injections, EEG/EMG and optical fiber implantation

For virus injections, a craniotomy (1 mm diameter) was drilled on top of the target site. Injections were performed using Nanoject II (Drummond Scientific) with pulled glass pipettes. The injection settings were 40 nL per pulse at the rate of 23 nL/s with 20 s interval between pulses. A mixture of equal amount of AAV-hSyn-FLEX-ChrimsonR-tdT (5.7× 10^12^ gc/mL, from University of North Carolina at Chapel Hill virus core) and AAV9-hSyn-GRAB_NE3.1_ (2-9 × 10^13^ gc/mL, WZ Biosciences) were injected bilaterally into the LC at multiple depths (-5.4 AP, ±0.9 ML, -3.4, -3.6 and -3.8 DV). For fiber photometry imaging at the mPFC, AAV9-hSyn-GRAB_NE3.1_ was injected (+1.8 AP, +0.3 ML, -2.0, -2.25 and -2.5 DV with 100nl at each depth).

For EEG/EMG recording, a reference screw was inserted into the skull on top of the left cerebellum. Two EEG electrodes were implanted about 3 mm posterior to bregma and 2.5 mm from the midline. Two EMG electrodes were inserted into the neck muscles. Insulated leads from the EEG and EMG electrodes were soldered to a pin header, which was secured to the skull using dental cement.

Fiber optic cannula (1.25-mm ferrule, 200-µm core) were implanted at -5.4 AP, ±0.9 ML, -3.5DV in the LC; and +1.8 AP, +0.3 ML and -2.0 DV in the mPFC. All screws, optical fibers and connectors required for EEG/EMG recordings were then secured onto the skull using dental cement. Animals were monitored till regaining motor ability before returning to the housing room.

### Close-loop sleep deprivation

Mice in their home cages were placed on an orbital shaker for habituation the day before sleep deprivation. An Internet-of-Things relay device (Digital Loggers) was used to control motion (∼200 revolutions per minute) of an orbital shaker (MT-201-BD, Labnique) with a TTL pulse from a RZ10X Processor (Tucker-Davis Technologies, TDT). For SD (ZT 1-4), a custom-written MATLAB program was used to calculate the real-time EMG activity of the mouse and generate TTL pulses to activate the shaker when EMG drops below a preset threshold indicative of sleep . Shaker activities introduced occasional sharp artifacts in the photometry data but rarely affected the calculation of laser-evoked NE responses. We removed the electric noise introduced by the shaker for displaying examples of EEM and EMG data in Fig. 1d and Fig. 3c; all other calculations were unfiltered.

### Polysomnographic recording and brain state classification

EEG/EMG signals were acquired using a TDT PZ5 amplifier and Synapse software, with a bandpass filter of 0.5–300 Hz and sampling rate at 1017 Hz. Spectral analysis was carried out using fast Fourier transform, and brain states were classified as previously described (wake: desynchronized EEG and high EMG activity; NREM: synchronized EEG with high-amplitude, low-frequency (1–4 Hz) activity and low EMG activity; REM: high EEG power at theta frequencies (6–9 Hz) and low EMG activity). The classification was made with 5-s bins using a custom-written graphical user interface (programmed in MATLAB, MathWorks) ^1^. Data were presented as the mean across animals in all plots.

### Optogenetic stimulation and Fiber photometry

To enable simultaneous laser stimulation and fiber photometry imaging, pulses generated from 635 nm red laser diode (RWD Life Science) were sent to the same dichroic mini-cube (Doric lenses) used for fiber photometry. Laser powers were titrated as follows as measured at the tip of the patch cables: 2 mW for unilateral LC activation (Fig. 2e-g); 4-6 mW for all other experiments. Test laser pulses had a width of 50 ms at random intervals between 80 – 160 s for the unilateral LC activation experiment (Fig. 2e-g), 40 - 80 s for all other experiments. For minute-scale LC activation experiments (Fig. 4d,e, and Extended Data Fig. 5), the inter-stimulation interval was 16 min.

Fiber photometry recording was performed using TDT RZ10x real-time processor. Fluorescence elicited by 405 nm and 465 nm LEDs (at 210 and 330 Hz, respectively) were filtered through the dichroic mini-cube (Doric lenses) and collected with an integrated photosensor. Signals were demodulated and pre-processed using the TDT Synapse software, with which optogenetic stimulation, fiber photometry and EEG/EMG recordings were synchronized.

### Calculation of laser-evoked NE release

For 6 h recordings (Fig. 1 to 3), 465nm signals for each session were z-scored and normalized to the mean of the first hour (ZT 0-1). Evoked NE responses for each pulse were calculated as the difference between 1s after laser and 1s before and binned every 6 min. Single exponential curve fitting was done in MATLAB with the formula: 1-b*exp(-x/t). For LC activation on minute scales (Fig. 4 and Extended Data Fig. 5), laser-evoked NE responses in NREM were normalized to the mean of 5 min before laser and binned every 40 s. Data were presented as the mean across animals in all plots.

### Calculation of laser-evoked brain states changes

Laser-evoked wakefulness was calculated as the percentage of wakefulness 20 s after laser minus that of 20 s before, binned every 15 min, and averaged across animals.

### Brain-state-dependent analysis

For brain-state-specific analysis in Fig. 4, we defined NREM to wake transitions as >30s of NREM followed by >30s of wakefulness. Similarly, wake to NREM transitions were defined as >30s of wakefulness followed by >30s of NREM. Wake bouts were selected for analysis if 85% wake were present in the following 12 min after transition; NREM bouts were selected if there was <15% wakefulness 12 min after wake to NREM transition. Short and long wake bouts were separated based on the median duration of all wake bouts (55 s). Laser-evoked NE responses were then binned every 30 s and averaged across all trials. Single exponential curve fitting was done in MATLAB with the formula: a-b*exp(-x/t). Fig. 4e was binned every 40 s and average across animals.

### Histology and immunohistochemistry

Mice were deeply anesthetized using isoflurane and transcardially perfused using PBS followed by 4% paraformaldehyde in PBS (Gibco). Brains were post-fixed overnight in 4% paraformaldehyde (Electron Microscopy Sciences), then de-hydrated overnight in a 30% sucrose PBS solution before cryopreservation in a -80°C freezer. Brains were embedded and mounted with Tissue-Tek OCT compound (Sakura Finetek) and sectioned in coronal positions at 30 μm using a cryostat (Thermo Scientific). For immunostaining, floating brain slices were washed 3 times with 1x PBS for 10 min each, permeabilized, and incubated in a blocking solution (5% normal bovine serum in 1x PBS, 0.4% Triton X-100) with primary antibodies overnight at 4 °C on a rotating shaker. The next day, after washing 3 times in 1x PBS for 10 min each, brain slices were incubated in fluorescently conjugated secondary antibodies (1:500, Invitrogen) for 2 h at room temperature. Finally, brain slices were washed in 1X PBS, counterstained with 4’,6-diamidino-2-phenylindole dihydrochloride (DAPI; Sigma-Aldrich) and mounted on slides with VECTASHIELD antifade Mounting Medium (Vector Laboratories, H-1000). Images were acquired using a fluorescence microscope (Keyence BZX-710) with the same exposure settings for WT and *Rptor*-/- pairs. For TH staining, each ROI (cell) was manually identified, and fluorescent signals were quantified using ImageJ. In total, 670 and 679 cells from 6 pairs of animals were quantified for *Rptor*-/- and WT animals, respectively.

Primary and secondary antibodies used are as follows:

Rabbit anti-TH antibody (ab112, abcam, 1:300)
Donkey anti-rabbit secondary antibody, Alexa fluorophore 647 (A-31573, Invitrogen; 1:500)

### Statistics

Statistical analysis was performed in MATLAB (paired or unpaired t-test) and R (ANOVA). Significance was determined at P < 0.05. For two-way mixed model ANOVA, random slopes and intercepts models were specified and mouse/subject were included as a random covariate using the *lme4* package^2^. Post hoc analyses were t-tests with Bonferroni corrections for multiple comparisons and were conducted using the *lsmeans* package^3^ in R. All statistical comparisons are two-sided and conducted on animal averages (i.e., each animal has one observation per level(s) of the independent variable).

---

**Extended Data Fig. 1.**
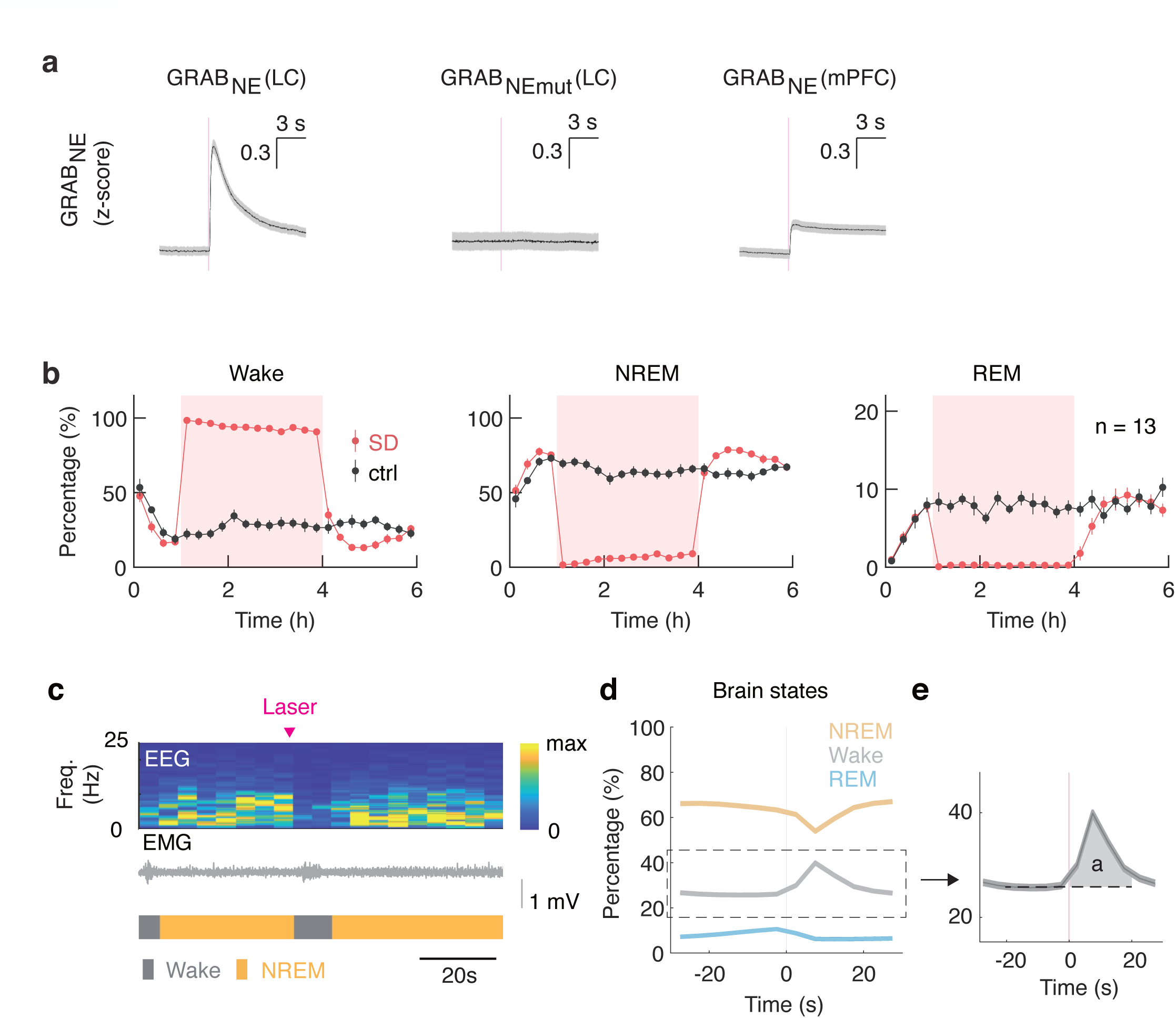
Laser-evoked NE release and brain state changes in SD experiment. **a,** Examples of laser-triggered GRAB_NE3.1_ fluorescence (z-score) in the LC and ipsilateral mPFC in one recording session. NE mutant sensor (GRAB_NEmut_) shows no laser induced responses. Shading, SEM. **b,** Percentages of each brain state binned every 20 min and averaged across animals in SD (red) or control (black) condition. **c,** An example showing that a single 50 ms pulse induced transient wakefulness as characterized by EEG activation and increased EMG activity. **d,** Percentages of each brain state before and after 50 ms laser pulses (average across mice, n = 13). **e,** Laser-evoked wakefulness is calculated as the difference in the percentage of wakefulness 20 s before and after laser.

**Extended Data Fig. 2.**
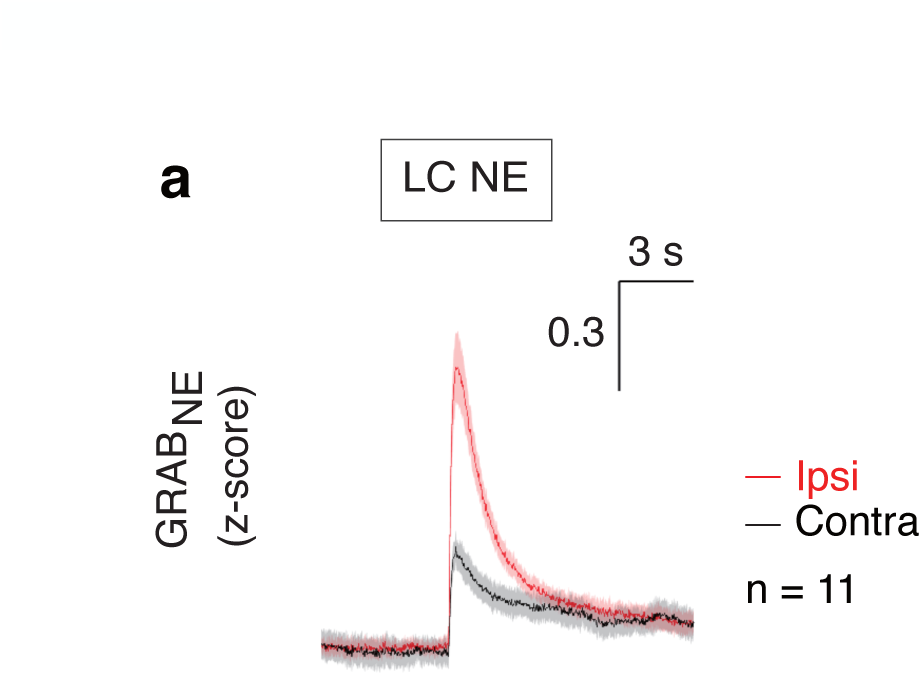
Crosstalk between the two sides in the LC activation experiment. **a,** Laser-evoked NE release on the activated side were ∼ 3-fold greater than that of the contralateral side (Average across mice, n =11). Shading, SEM.

**Extended Data Fig. 3.**
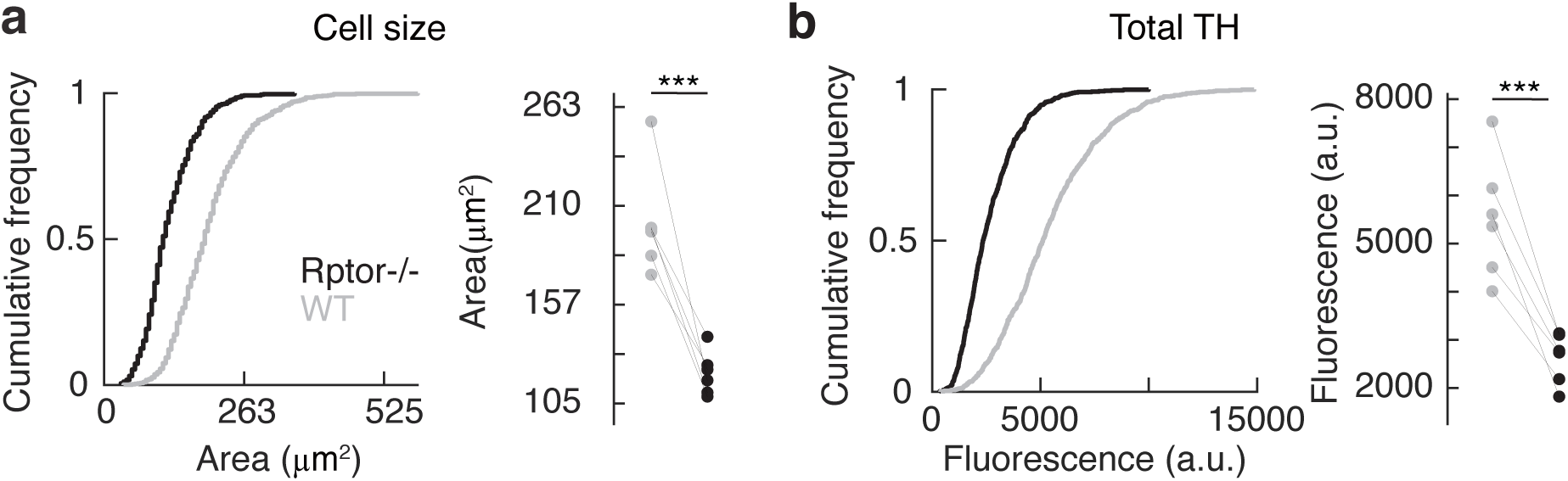
Reduced LC neuron cell size and TH abundance in *Rptor*-/- mutants. **a&b,** Cumulative frequency of cell size (**a**) and total TH fluorescence (**b**) of individual LC neurons in *Rptor*-/- (n = 6) and WT (n = 6) mice. P_Cell size_ = 0.002; P_total TH_ = 0.001 (paired t-test).

**Extended Data Fig. 4.**
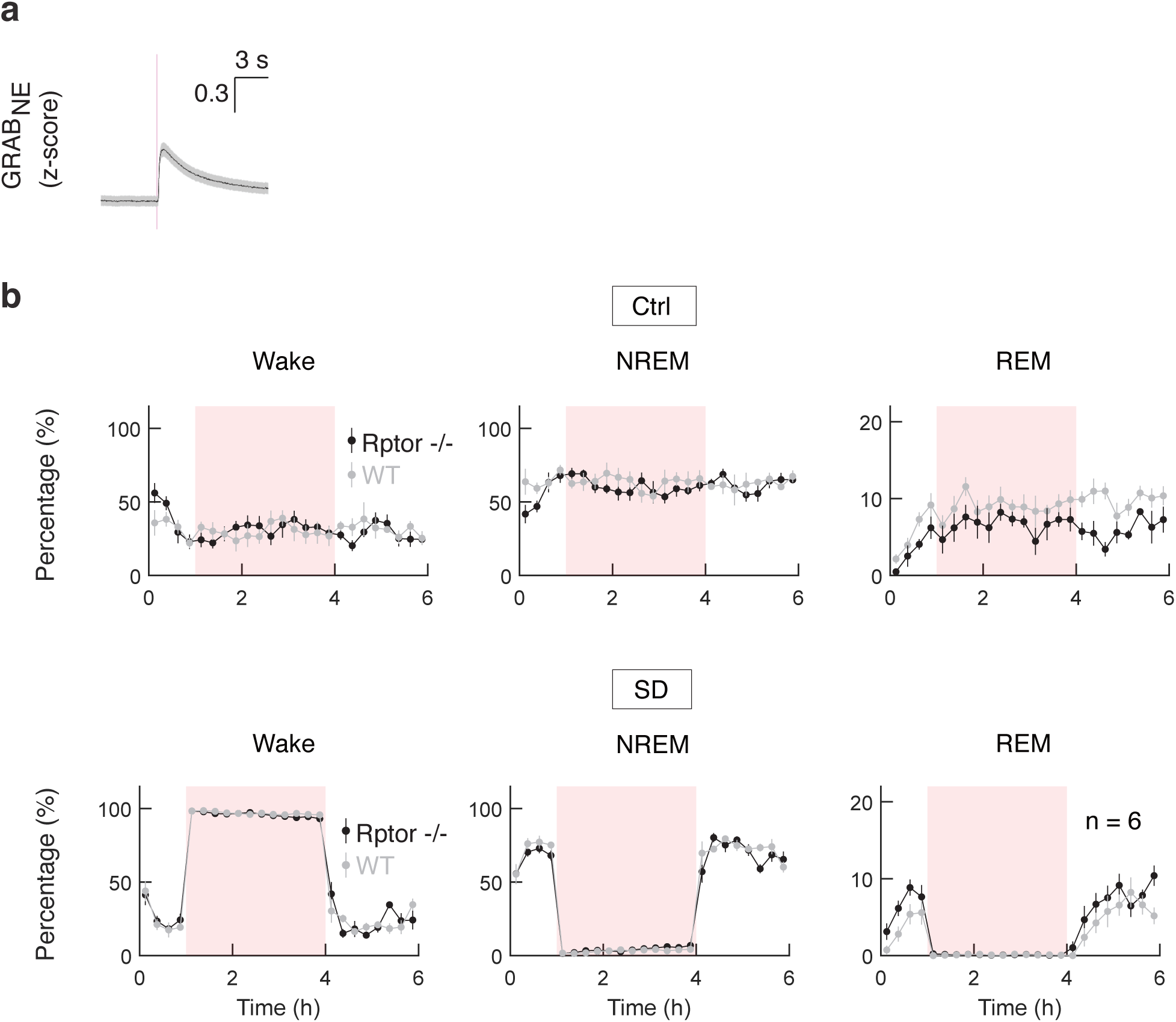
Brain states in the SD experiment of *Rptor*-/- mutants. **a,** 50 ms laser pulses elicited robust NE response in the LC in *Rptor*-/- mutants, although with a smaller peak amplitude than WT. Plotted example from one recording session. Shading, SEM. **b,** Percentages of each brain state averaged across mice for *Rptor*-/- (black, n = 6) or WT (grey, n = 6) in control (upper panel) or SD (lower panel) condition. Data points are binned every 20 min.

**Extended Data Fig. 5.**
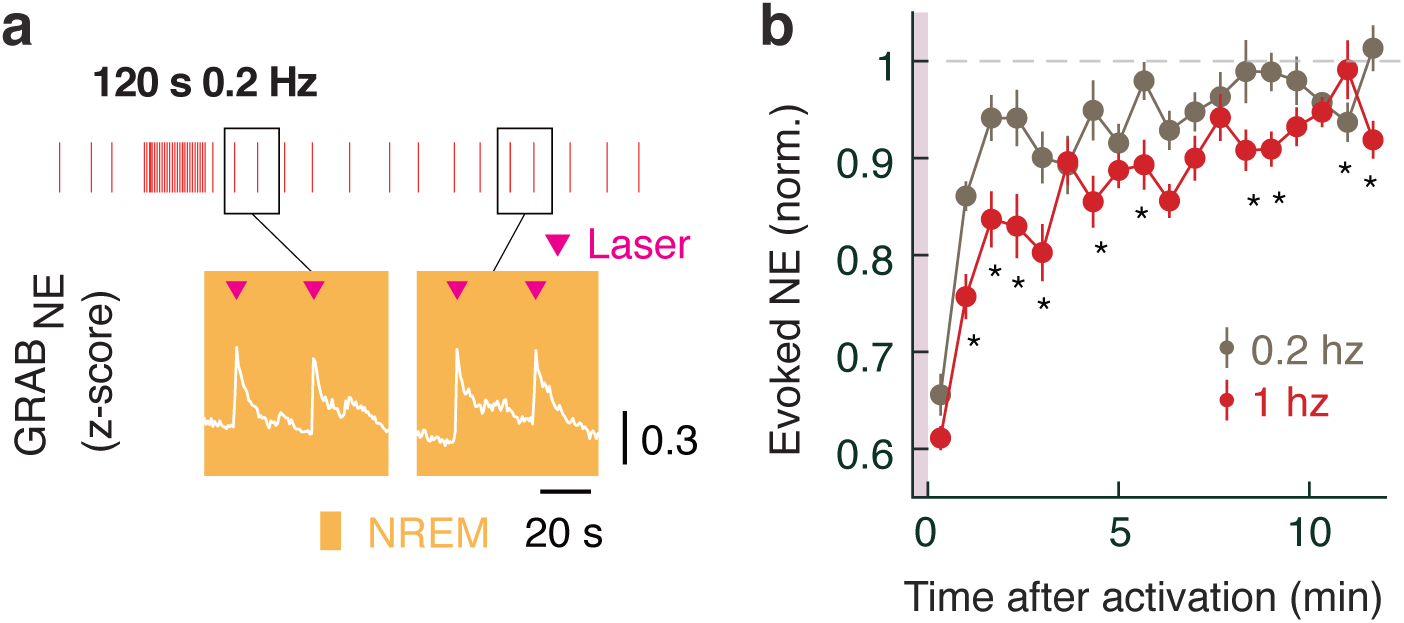
Recovery rate of LC NE transmission depends on frequency of LC activation. **a,** Example GRAB_NE3.1_ fluorescence showing evoked NE release in the LC after 0.2 hz LC activation for 120s. Vertical bars represent 50 ms laser pulses. **b,** Laser-evoked NE response in NREM after LC activation at the frequency of 0.2 (grey, n = 6) or 1 Hz (red, n = 7). Data points are averaged across animals and binned every 40 s. P = 0.009, P_treatment_ = 1 × 10^−11^, P_time_ < 2.2 × 10^−16^; * P < 0.05 (two-way ANOVA with Bonferroni correction).

## Notes

### Competing Interest Statement

The authors have declared no competing interest.

